# Comparing transcriptional dynamics of the epithelial-mesenchymal transition

**DOI:** 10.1101/732412

**Authors:** David P. Cook, Barbara C. Vanderhyden

**Affiliations:** Cancer Therapeutics Program, Ottawa Hospital Research Institute, Ottawa, ON, Canada; Department of Cellular and Molecular Medicine, University of Ottawa, Ottawa, ON, Canada

## Abstract

Epithelial-mesenchymal (E/M) heterogeneity is ubiquitous within all epithelial tissues and the reversible transition between these two states provides cells with plasticity that contributes to organogenesis in the developing embryo, tissue homeostasis in adults, and tumour progression^1^. While the epithelial-mesenchymal transition (EMT) has been extensively studied, no common, EMT-defining gene expression program has been identified^2^. Here, we leverage highly multiplexed single-cell RNA sequencing (scRNA-seq) to compare the transcriptional program associated with the EMT across a variety of contexts, assessing 103,999 cells from 960 samples, comprising 12 EMT time course experiments and 16 independent kinase inhibitor screens. We demonstrate that the EMT is not simply a linear transition between E/M states, and transcriptional dynamics are widely variable across contexts, regardless of the cell type and cytokine used to induce the transition. While many canonical EMT genes were poor markers of the transition in our models, we identified 86 conserved mesenchymal-associated genes also coexpressed in a variety of mouse and human epithelial and carcinoma tissues. Despite the heterogeneous transcriptional responses, we identified a core set of largely conserved transcription factors coordinating these dynamics, including RELB and SOX4. Finally, we found that the EMT is associated with a broad increase in expression of secreted factors. Kinase inhibitor screens revealed multiple paracrine dependencies of the EMT, including a novel association between TGFB1 and the TNF-associated kinase RIPK1. Together, these results comprehensively highlight the complexity and diversity of the EMT, but also reveal dynamics conserved across contexts. This work will provide the foundation for understanding the nature of E/M heterogeneity and its functional consequences, which could elucidate various physiological processes and be leveraged for cancer treatments.

---

To assess transcriptional dynamics of the EMT across a variety of contexts, we used MULTI-seq^3^ to generate single-cell RNA sequencing (scRNA-seq) data from twelve distinct EMT time course experiments. We assessed four different cancer cell lines capable of undergoing an EMT (A549, lung; DU145, prostate; MCF7, breast; OVCA420, ovarian) and exposed each to known EMT-inducing factors: TGFB1, EGF, and TNF **(Extended Data Fig. 1a)**. For each condition, samples were collected at eight distinct time points, including three after the EMT-inducing stimulus had been removed **(Fig. 1a)**. In the aggregated data, expression profiles clustered dominantly by cell line, and after demultiplexing, the majority of cell line annotations (95.8% on average) were restricted to a dominant cluster, demonstrating robust multiplexing **(Fig. 1b, Extended Data Fig. 1b)**. In total, we annotated 58,088 single cells from across 576 samples, comprising six replicates of the twelve time course experiments **(Fig. 1c, Extended Data Fig. 1c)**. Replicates were highly correlated, supporting the consistency of the experimental procedures and processing workflow **(Extended Data Fig. 2**).

**Figure 1.**
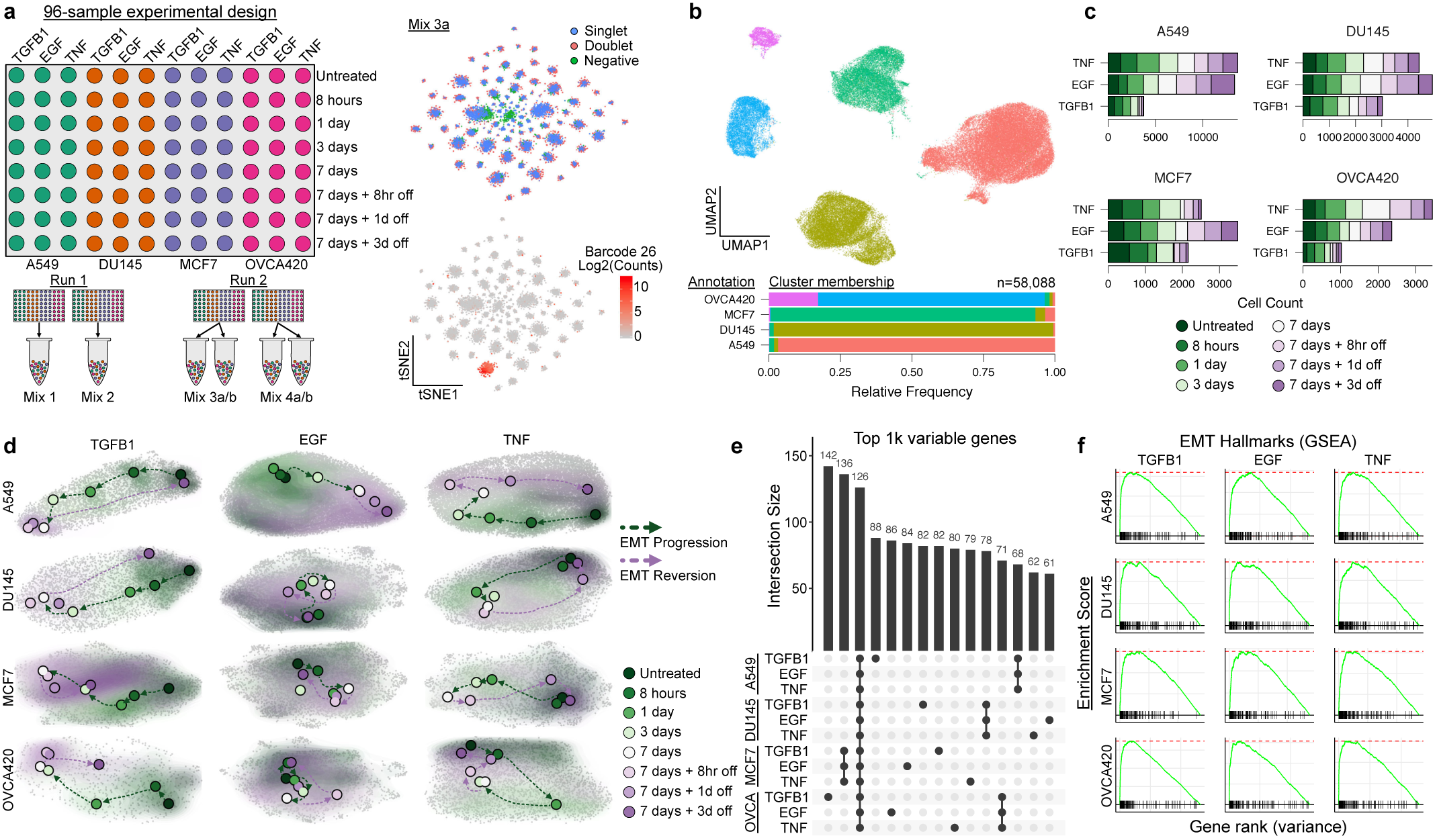
Multiplexed scRNA-seq enables transcriptional profiling of twelve EMT time course experiments. **a**, Schematic of the 96-well experimental design for the 12 EMT time course experiments (left), and t-SNE embeddings of the MULTI-seq barcode counts, demonstrating strong signal for demultiplexing (right). **b**, UMAP embedding of aggregated expression data of all data, coloured by unsupervised clustering (top), and a graph showing the relative proportion of annotations for each cell line assigned to each cluster after demultiplexing (bottom). **c**, Graph showing the number of cells captured for each time course experiment. **d**, UMAP embeddings of each of the 12 time course experiments. Grey dots correspond to individual cells, shaded regions represent the related sample density for each timepoint, and coloured dots correspond to the maxima of the density function. **e**, UpSet plot showing the intersections of the top 1000 variable genes of each time course experiment. **f**, GSEA showing the enrichment of the MSigDB EMT Hallmark gene set in top variable genes from A549 cells treated with TGFB1 (top), and graphs showing the normalized enrichment scores (NES) for all conditions.

We next isolated data for each of the twelve time course experiments to assess the temporal progression. In each case, time-dependent shifts in cells’ expression profiles were evident **(Fig. 1d)**. While the top 1000 variable genes for each time course showed expression patterns conserved across cell lines, context-dependent gene sets were also prevalent **(Fig. 1e)**. Gene set enrichment analysis (GSEA) on variance-ranked genes for each time course, however, did demonstrate consistent enrichment for EMT-associated expression **(Fig. 1f)**.

To specifically compare temporal dynamics of the EMT across contexts, we first pseudotemporally ordered the cells from each condition **(Fig. 2a, b)**. In each time course, cells progressively transitioned for the full 7 days of EMT induction, and withdrawal of the EMT stimulus led to a near complete reversion after as few as three days **(Fig. 2b)**. While interesting that the rate of reversion was faster than EMT progression itself, the rapid removal of paracrine factors when changing culture media likely accelerated the process.

**Figure 2.**
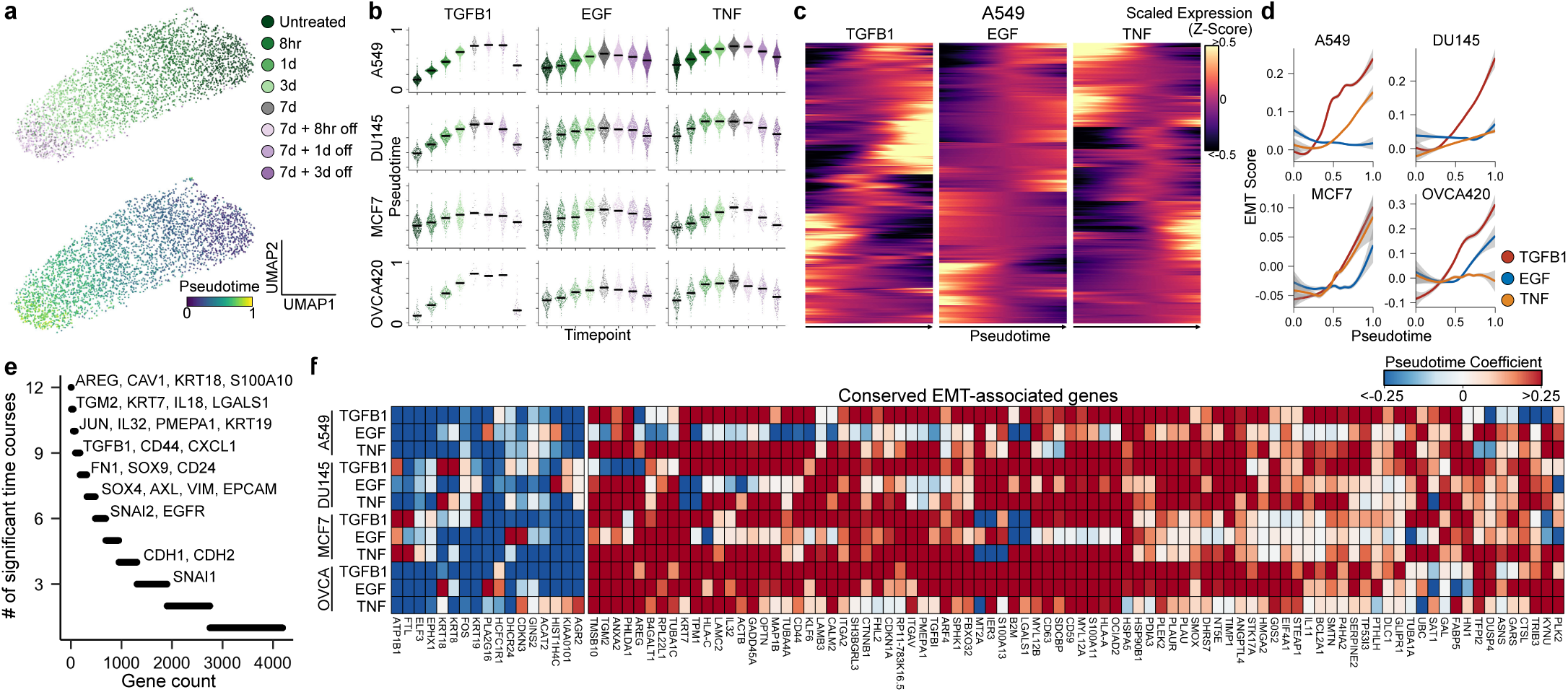
Comparing transcriptional dynamics of the EMT reveals global expression patterns are remarkably context specific. **a**, UMAP embeddings of A549 cells treated with TGFB1. Each point represents an individual cell and colours correspond to time point (top) or pseudotime value (bottom). **b**, Sina plot showing the distribution of pseudotime values across timepoints for all 12 time course experiments, with timepoints coloured the same as in (a). **c**, Expression dynamics of differentially expressed genes throughout pseudotime for A549 cells treated with TGFB1, EGF, or TNF. **d**, Smoothed model of the EMT hallmark gene set score throughout pseudotime. **e**, Counts of how frequently each gene is differentially expressed among time course experiments. **f**, Heatmap showing EMT-associated expression changes associated with a gene set of all genes that are differentially expressed in at least 8 time course experiments. The colormap corresponds to the pseudotime beta coefficient of the linear model for each gene.

We next assessed gene expression dynamics throughout the pseudotemporal trajectories. In all cases, transitions were not simply linear processes of two inversely related epithelial and mesenchymal expression programs. Rather, all transitions involved combinations of transcriptional events **(Fig. 2c, Extended Data Fig. 3a)**, suggesting that the EMT may be a multi-step progression. We found that each condition, with the exception of A549 cells induced with EGF and OVCA420 treated with TNF, was associated with an average increase in the expression of the MSigDB Hallmark EMT gene set^4^, with TGFB1 often producing the most potent effects **(Fig. 2d)**. GSEA revealed, however, that differentially expressed genes from these two conditions, along with all others, were enriched for the same gene set **(Extended Data Fig. 3b)**. Assessing specific expression dynamics of the gene set, it is evident that the net-neutral EMT score for these data sets is due to some EMT genes being upregulated, while others are downregulated **(Extended Data Fig. 3c)**.

Surprisingly, of all genes differentially expressed across conditions, the majority changed in as few as 1-2 conditions, suggesting that the global expression programs associated with the EMT are remarkably context-specific **(Fig. 2e)**. A small subset of canonical EMT genes, including TGFB1, CD44, and FN1, along with less-reported genes, such as TGM2 and PMEPA1, were differentially expressed in most conditions. Interestingly, expression of other canonical genes, including SNAI1, CDH1 (E-cadherin), and CDH2 (N-cadherin) was only changed in a small number of conditions **(Fig. 2e)**. In fact, the majority of the MSigDB Hallmark EMT gene set was differentially expressed in only a small number of conditions, with only 49/200 hallmark genes being differentially expressed across the majority of conditions **(Extended Data Fig. 4a)**. To identify signatures that may not have been represented in the hallmark gene set, we took all genes that were differentially expressed in at least 8 of our experimental conditions and compiled our own geneset of 86 conserved upregulated genes and 17 downregulated genes **(Fig. 2f)**. We also confirmed that these 86 EMT-associated genes are enriched in the top variable genes of cancer cells from human lung tumours and syngeneic mouse tumour models, as well as healthy epithelium from various mouse tissues **(Extended Data Fig. 4b)**. Further, in each of these data sets, the 86 EMT-associated genes are highly correlated **(Extended Data Fig. 4c)**, supporting that this gene set is coexpressed in vivo and may be a reliable signature of the mesenchymal state in a variety of contexts.

While global transcriptional programs are poorly conserved across conditions, we hypothesized that conserved regulatory systems may exist, but give rise to diverse transcriptional output due to factors such as the cells’ epigenetic landscape. Across the experimental conditions we assessed, canonical EMT transcription factors—other than SNAI2—were rarely differentially expressed **(Fig. 3a)**. To identify conserved transcription factors that may be contributing to the transition, we scored each cell for the coexpression of transcription factors and their putative target genes (regulons) and identified those that showed differential activity throughout the EMT (Fig. 3b). Throughout the EMT, RELB, ATF4, SOX4, and KLF6 regulons showed frequent activation, whereas ELF3 and MYBL2 activity decreased **(Fig. 3c)**. These factors have all been previously implicated in the EMT, but are not typically considered core EMT regulators^5–10^. To validate these results, we performed ATAC-seq on a TGFB1 time course experiment in OVCA420 cells and assessed the accessibility of transcription factor motifs throughout the transition **(Fig. 3d)**. In many cases, including RELB, we found that motif accessibility mirrored regulon activity measured from scRNA-seq data alone **(Fig. 3e)**.

**Figure 3.**
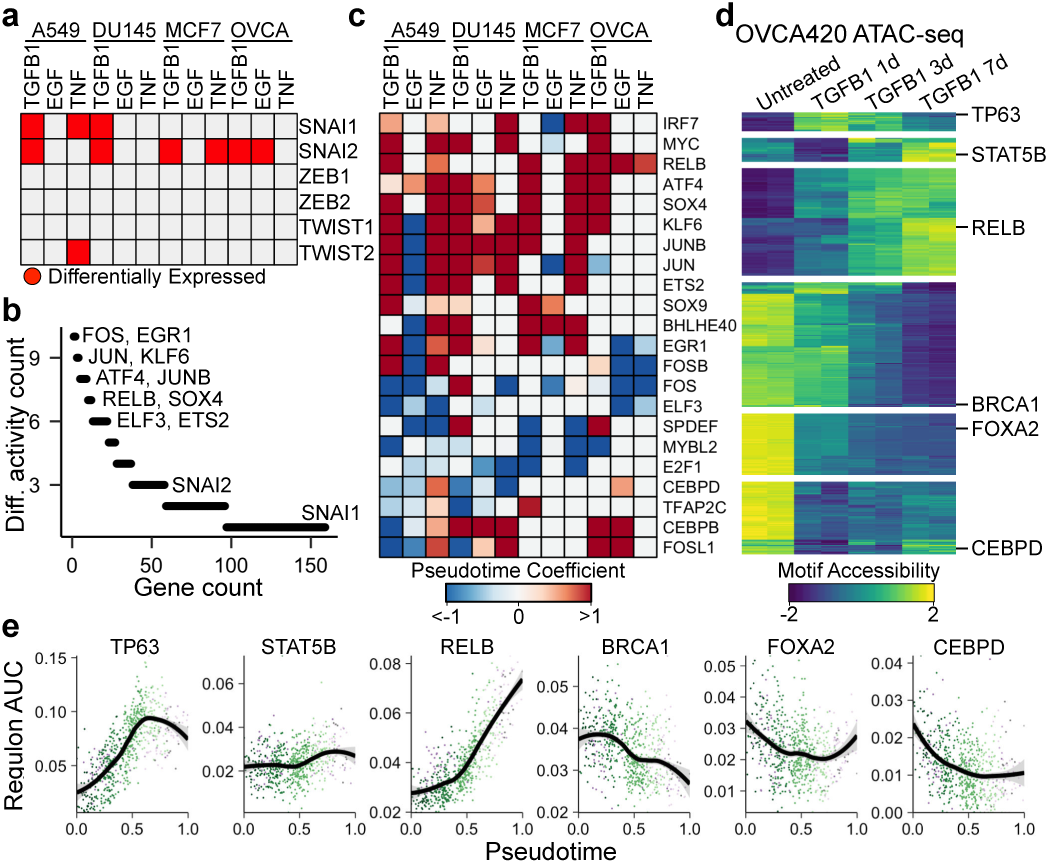
Inferring transcription factor activity throughout the EMT. **a**, Plot showing in which time course experiments various canonical EMT transcription factors are differentially expressed. **b**, Counts of how frequently various transcription factors and their associated regulons are differentially active among time course experiments. **c**, Heatmap showing EMT-associated changes of regulons that are differentially active in at least 6 time course experiments. The colormap corresponds to the pseudotime beta coefficient of a linear model for each regulon. **d**, Differential accessibility of transcription factor motifs from ATAC-seq data of OVCA420 cells treated with TGFB1 for 0, 1, 3, or 7 days. The colormap represents the accessibility Z-score for each transcription factor motif. Examples of transcription factors from each cluster are listed. **e**, Regulon activity score of the same transcription factors listed in (d) inferred from the OVCA420 TGFB1 time course experiment. Each dot represents a single cell, coloured by time point. The black line corresponds to the modelled trend from a generalized additive model.

Cytokines and various growth factors were prominent among lists of differentially expressed genes in each condition. In fact, we found that total average expression of secreted factors spanning a variety of signalling pathways increased in each of our 12 time courses **(Fig. 4a, b)**. This supports previous observations about the involvement of paracrine signalling during the EMT^11^, which could both coordinate the transition across a population of epithelial cells, and modulate the cells’ microenvironment (eg. immunosuppression in cancer)^12–14^. Given this, we next established an experimental design to mechanistically assess the dependence of the EMT on paracrine signalling. We curated a selection of 22 small molecule inhibitors and treated cell lines alone for 7 days, or in combination with one of the three EMT inducers previously used **(Fig. 4c)**. Leveraging MULTI-seq to multiplex samples, we ultimately generated scRNA-seq profiles for 45,911 cells across the 384 distinct conditions.

**Figure 4.**
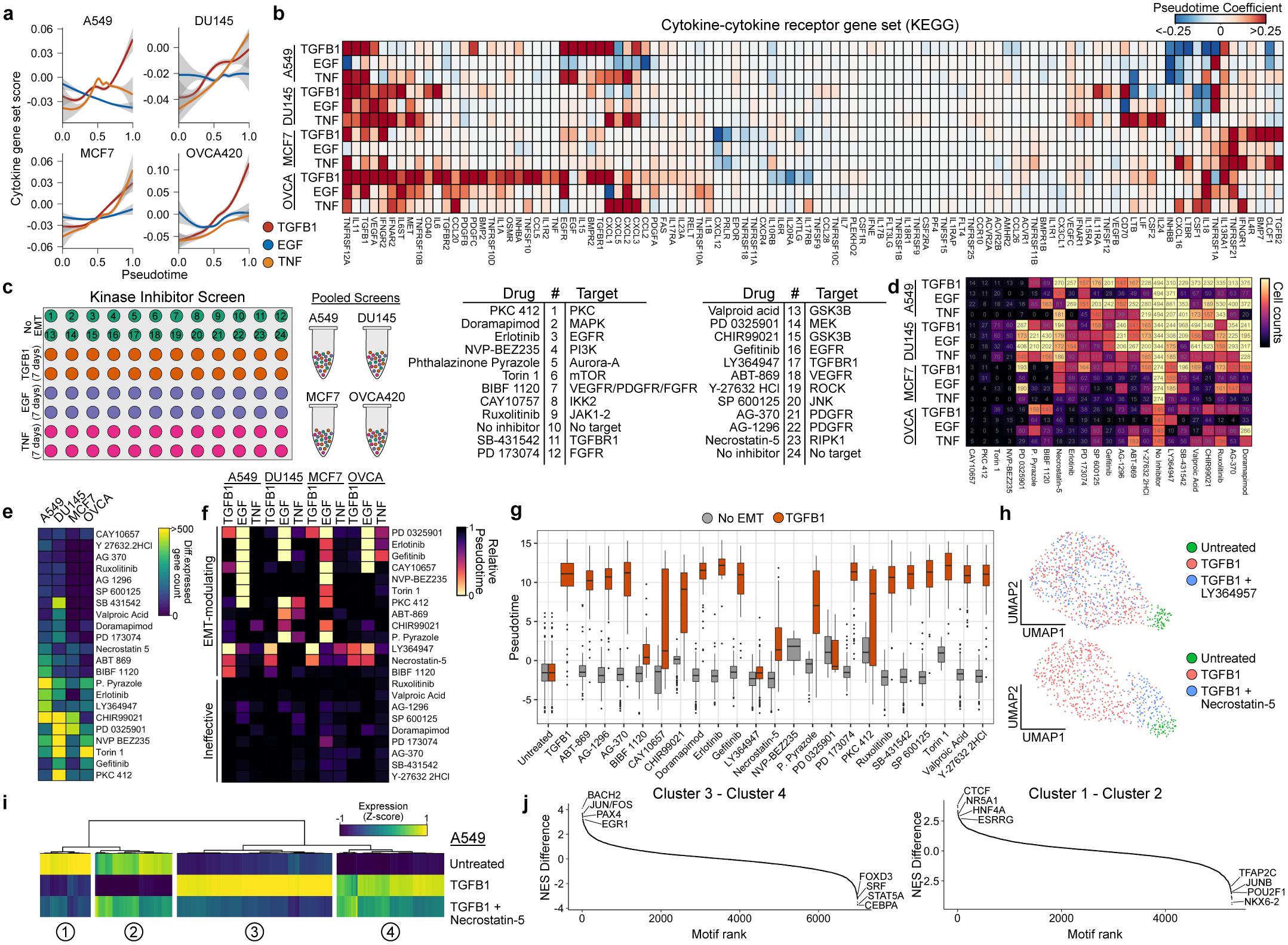
Kinase inhibitor screens demonstrate the dependence of the EMT on paracrine signalling. **a**, Gene set score of the KEGG pathway “Cytokine-cytokine receptor interaction” over pseudotime for each time course experiment. **b**, Heatmap showing EMT-associated changes of the individual genes of the same gene set as in (a), only listing those with a significant change in at least one time course experiment. **c**, Schematic of the 384-sample experimental design for the kinase inhibitor screen. **d**, Heatmap showing the number of cells annotated to each condition after demultiplexing the scRNA-seq data. **e**, Summary of the number of genes that are differentially expressed in each cell line exposed to the inhibitors without EMT induction. **f**, Average pseudotime values calculated for each condition. **g**, Boxplots showing the distribution of pseudotime values for A549 cells treated with the inhibitors alone (grey) or in combination with TGFB1 (orange). **h**, UMAP embeddings of untreated A549 cells with those had been treated with TGFB1 alone or in combination with the TGFBR1 inhibitor LY364947 (top), or the RIPK1 inhibitor Necrostatin-5 (left). **i**, Heatmap showing expression (Z-score) of genes differentially expressed in A549 cells by TGFB1 in untreated A549 cells, as well as those treated with TGFB1 alone or in combination with Necrostatin-5. **j**, Difference in normalized enrichment scores for transcription factor targets in genes that are successfully inhibited by Necrostatin-5 compared to those that are not. Positive values correspond to regulons with that are enriched in Necrostatin-5-inhibited genes, whereas negative values represent those are not affected by Necrostatin-5.

From retrieved cell counts alone, drop-out patterns of cell line-dependent and -independent cytotoxic/cytostatic effects can be observed (Fig. 4d). For example, PKC 412 (PKC), NVP-BEZ235 (PI3K), CAY10657 (IKK2), and Torin 1 (mTOR) largely affected cell counts of all cell lines at the time of processing, and the cells that were captured had dramatic changes to their expression profiles **(Fig. 4e, Extended Data Fig. 5)**. PD 0325901 (MEK) had cytotoxic/cytostatic effects on A549 and OVCA420 cells, but not DU145 or MCF7 cells, and yet, all cells treated with the inhibitor had a large number of differentially expressed genes **(Fig. 4d, e, Extended Data Fig. 5)**. CHIR99021 (GSK3B), in contrast, had little effect on cell viability but large effects on gene expression in all cell lines other than OVCA420 **(Fig. 4e)**.

To assess the impact of these inhibitors on EMT progression, we calculated pseudotime values for the inhibited cells using the models built from the corresponding time course experiment of the same cell line and EMT inducer **(Fig. 4f)**. From this, we could identify inhibitors that reduced cells’ pseudotime values at 7 days compared to uninhibited controls, therefore dampening the EMT response. LY364947 (TGFBR1), for example, abrogated TGFB1-induced EMTs **(Fig. 4f, g, h)**, and Erlotinib and Gefitinib (both EGFR) consistently blocked the effects of EGF **(Fig. 4f)**. The effects of these inhibitors, however, weren’t limited to directly blocking the EMT-inducing factor. For example, TGFBR1 inhibition partially blocked EMT progression in EGF-treated A549 and OVCA420 cells, and TNF-treated A549 and MCF7 cells **(Fig. 4f)**. MEK inhibition (PD 0325901), while having basal effects on untreated cells, prevented EMT progression in a variety of contexts, and the triple inhibitor of VEGFR, PDGFR, and FGFR (BIBF 1120) abrogated the effects of TGFB1 in A549 and DU145 cells **(Fig. 4f)**.

Inhibition of RIPK1—a kinase involved in activating NFKB and necroptosis pathways—with Necrostatin-5 prevented EMT progression in all of the same conditions as TGFBR1 inhibition, but was less potent in all cases **(Fig. 4f, g)**. Interestingly, RIPK1 inhibition only abrogated a subset of TGFB1-induced expression changes in each case, blocking the cells’ progression part way through the EMT **(Fig. 4h, i)**. Exploring the expression program of RIPK1-inhibited cells, we found that the inhibitor largely prevented dynamics associated with BACH2, SMARCC1, and JUN/FOS target genes, whereas uninhibited genes were enriched for CEBPA, STAT5A, and SRF targets **(Fig. 4j)**.

Here, we have highlighted the complexity and diversity of the EMT, demonstrating that the majority of transcriptional changes associated with the transition are context-specific. However, while the molecular basis of the EMT may be diverse, it will be important for future studies to determine the impact of this variation on cellular function. We have identified a small, but largely conserved group of genes associated with E/M heterogeneity both in our experimental systems and in vivo. Given the inconsistencies in canonical EMT-associated expression patterns, we suggest that these conserved genes may be more reliable for assessing E/M in cells. This study, however, raises important questions about whether the notion of “E/M status” is appropriate at all. While the EMT is often conceptualized as a single linear continuum, our data support a model where the transition is a multidimensional nonlinear process, dependent on the combination of factors driving the transition and the epigenetic state of the cells. Finally, we also provide evidence to support the involvement of paracrine signalling in the EMT, identifying novel interactions between signalling pathways, such as TGFB1 and RIPK1. Given this, properties of the microenvironment are also likely to modulate the transition, promoting further complexity. Ultimately, untangling this complexity and its association with cellular function will be critical to fully understand the involvement of the EMT in a variety of biological processes.

## Methods

### Cell culture

A549, DU145, and MCF7 cells were obtained from ATCC (CCL-185, HTB-81, and HTB-22, respectively). OVCA420 cells were kindly provided by Dr. Gordon Mills. All cells were cultured in Dulbecco’s Modified Eagle Medium (DMEM) with 4.5g/L glucose, L-glutamine, and sodium pyruvate (Corning, 10-013-CV), supplemented with 10% of fetal bovine serum (FBS) and cultured at 37°C with 5% CO2.

### EMT time course experiments

For each cell line, 10,000 cells were plated into each well of a 96-well plate according to the schematic in Figure 1a. The addition of TGFB1, EGF, and TNF were scheduled such that all timepoints completed at the same time for collection. Cells were treated with 10ng/mL TGFB1 (R&D Systems, #240-B-010), 30ng/mL EGF (Invitrogen, #PHG0311), or 10ng/mL TNF (Invitrogen, #PHC3015). Media was changed and fresh TGFB1, EGF, or TNF were added every two days to ensure relatively constant concentrations of these factors. To avoid over-confluence throughout the experiments, cells were passaged as required, but not within the last two days of the time course to avoid artifacts at the time of collection. After the scheduled treatments, cells were immediately processed for scRNA-seq multiplexing.

The time course experiments were performed twice independently. Each time, the two time course replicates were performed in parallel, and on the second time through the experiment, two 10x libraries were generated for each plate replicate. Samples from the first replicate are labelled “Mix1” and “Mix2”, corresponding to the two plates running in parallel. Samples from the second replicate are labelled “Mix3a/b” and “Mix4a/b”.

### Kinase inhibitor screen

For each cell line, 10,000 cells were plated into four 96-well plates according to the schematic in Figure 4c. Cells were simultaneously treated with small molecule kinase inhibitors (listed in Figure 4c) and either 10ng/mL TGFB1, 30ng/mL EGF, or 10ng/mL of TNF. No-inhibitor and No-EMT controls were also included for all conditions. All inhibitors were used at a final concentration of 1µM (Cayman Chemical Kinase Screening Library, Item No. 10505, Batch No. 0537554). EMT inducers and kinase inhibitors were refreshed daily after replacing the culture media. After 7 days of treatment, all samples were immediately processed for scRNA-seq multiplexing.

### Multiplexing individual samples for scRNA-seq

Multiplexing was performed according to the MULTI-seq protocol^3^, and reagents were kindly provided by Dr. Zev Gartney. Briefly, culture media was removed and each well was washed with 1x Dulbecco’s Phosphate-Buffered Saline (PBS; Corning, #21-031-CV). Next, a lipid-modified DNA oligonucleotide (LMO) and a unique “sample barcode” oligonucleotide were added at 200nM to 0.05% trypsin with 0.53mM EDTA. This was added to each sample to be multiplexed, with each sample receiving a different sample barcode. Cells were incubated with this trypsin mixture for 5 minutes at 37°C, and plates were gently mixed periodically. After 5 minutes, a common lipid-modified co-anchor was added to each well at 200nM to stabilize the membrane residence of the barcodes. Cells were incubated for an additional 5 minutes at 37°C with periodic mixing. After this labelling time, all cells were in suspension, lifted from the plate. The trypsin was then neutralized with cultured media, and the cells were mixed by pipetting to ensure a single cell suspension. Samples were then transferred to V-bottom 96-well plates, and pelleted at 400xg for 5 minutes. Barcode-containing media was removed, and the cells were then washed with PBS + 1% bovine serum albumin (BSA). Washes were performed twice, and after the final wash, cells were resuspended in PBS + 1% BSA, pooled together, re-pelleted, and resuspended in PBS + 1% BSA. Viability and cell counts were then performed, before preparation of the scRNA-seq libraries.

### scRNA-seq library preparation and sequencing

Single-cell suspensions were processed using the 10x Genomics Single Cell 3’ RNA-seq kit (v2 for time course experiments, v3 for kinase inhibition). Gene expression libraries were prepared according to the manufacturer’s protocol. MULTI-seq barcode libraries were retrieved from the samples and libraries were prepared independently, as described previously^3^. Final libraries were sequenced on a NextSeq500 (Illumina). Expression libraries were sequenced so that time course libraries reached an approximate depth of 40,000-50,000 reads per cell (for the v2 scRNA-seq kit), and 20,000-25,000 reads per cell for the kinase inhibitor experiment (v3 scRNA-seq kit).

### Processing of raw sequencing reads

Raw sequencing reads from the gene expression libraries were processed using CellRanger v2.2.0 for the time course data, and v3.0.2 for the kinase inhibitor data. The GRCh38 build of the human genome was used for both. Except for explicitly setting --expect-cells=25000, default parameters were used for all samples. MULTI-seq barcode libraries were simply trimmed to 26bp (v2 kit) or 28bp (v3 kit) using Trimmomatic^18^ (v0.36) prior to demultiplexing.

### Demultiplexing expression data with MULTI-seq barcode libraries

Demultiplexing was performed using the deMULTIplex R package (v1.0.2) (https://github.com/chris-mcginnisucsf/MULTI-seq). The key concepts for demultiplexing are described in McGinnis et al.^3^. Briefly, the tool takes the barcode sequencing reads and counts the number of times each of the 96 barcodes appears for each cell. Then, for each barcode, it assesses the distribution of counts in cells and determines an optimal quantile threshold to deem a cell positive for a given barcode. Cells positive for more than one barcode are classified as doublets and are removed. Only cells positive for a single barcode are retained for downstream analysis. As each barcode corresponds to a specific sample in the experiment, the sample annotations can then be added to all cells in the data set.

### Data quality control and processing

Quality control was first performed independently on each 10x Genomic library, and all main processing steps were performed with Seurat v3.0.2^19^. Expression matrices for each sample were loaded into R as seurat objects, only retaining cells with more than 200 genes detected. Cells with a high percentage of mitochondrial gene expression were also removed. We then subsetted the data, making independent seurat objects for each time course or kinase inhibition experiment (ie. for all independent cell line and EMT inducer combinations). Each condition was then processed independently with a standard workflow. We first removed genes detected in fewer than 1% of the cells for the given experiment. The expression values were then normalized with standard library size scaling and log-transformation. The top 3000 variable genes were detected using the “vst” selection method in Seurat. Expression values were scaled and the following technical factors were regressed out: percentage of mitochondrial reads, number of RNA molecules detected, cycle cycle scores, and for the time course data, batch was also included. For initial exploration, PCA was run on the variable genes, but all UMAP embeddings included in figures are based on PCA run on genes used for pseudotemporal ordering of cells. UMAP embeddings were calculated from the first 30 principal components.

### Pseudotemporal ordering of cells

Pseudotime models for each time course experiment were built using the R package psupertime v0.2.1^20^ on the top 3000 variable genes from each condition. Psupertime is based on ordinal logistic regression, taking scRNA-seq data with sequential labels and identifying a linear combination of genes that places the cells in the specified label order. To build the pseudotime model for each time course, we first omitted the treatment withdrawal samples. Because psupertime is based on regression, however, pseudotime values for new data can be calculated by simply performing matrix multiplication between the coefficient matrix of the pseudotime model and the expression matrix of the new data. We used this approach to calculate pseudotime values for both the treatment withdrawal samples of the time course experiment. We also used the time course models to calculate pseudotime values for the respective kinase inhibition experiments. As the range of pseudotime values can vary between conditions, we simply rescaled the values from 0-1 in cases where multiple models were compared in the same figure.

### Differential expression analysis

For time course experiments, expression dynamics of each gene, or transcription factor regulon score, as a function of pseudotime was modelled using the generalized additive model function provided by the R package mgcv with the model exp ∼ s(pseudotime, k=4) + batch, with the smoothing parameter estimation method set to restricted maximum likelihood (method=”REML”). The number of basis functions (k) was chosen such that the residuals were randomly distributed. P-values associated with the smoothed pseudotime function for each gene were adjusted using the p.adjust() function in R with the Benjamini-Hochberg method. As many genes may significantly vary throughout pseudotime but have low effect sizes, we only evaluated significant genes (adjusted p-value < 0.05) that are also within the top 2000 variable genes of each time course experiment. While others may be biologically relevant, their signal in the data is often too low to assess reliably.

When assessing transcription factor activity (Fig. 3) and cytokine production (Fig. 4), we were more generally interested in assessing the “directionality” of change over pseudotime, so in these cases, we used the same approach, but removed the smoothing function from the model. This allowed us to report the single coefficient associated with the pseudotime covariate, representing whether activity generally increased or decreased throughout the transition.

For the kinase inhibition experiment, we assessed the number of differentially expressed genes in cell lines treated with a kinase inhibitor, but no EMT inducer. For this, we still used the gam() function provided by the mgcv package with the model exp ∼ inhibitor, setting the no-inhibitor controls as the intercept. We then quantified the number of genes with an adjusted p-value < 0.05.

### Calculating smoothed expression trends

To calculate smoothed expression trends over pseudotime, we used models used for differential expression, but calculated the fit values for 200 evenly-spaced pseudotime values ranging between the minimum and maximum pseudotime values.

### Gene set enrichment analysis

Gene set enrichment analysis (GSEA) was performed using the R package fgsea^21^. Input genes were ranked either by their variance values after the variance stabilizing transformation (“vst”), computed by Seurat’s FindVariableFeatures() function, or by adjusted p-value from the differential expression analysis. Reference gene sets were collected from the Molecular Signatures Database (MSigDB) v6.2.

### Gene set scoring

Gene set scoring of the EMT hallmark gene set and the KEGG pathway “Cytokine-cytokine receptor interaction” was performed using the AddModuleScore() function provided by the Seurat package. Default parameters were used.

### Transcription factor regulon scoring of single-cells

Regulon scores for individual cells were computed using the SCENIC workflow^22^. Log-transformed expression values for each time course experiment were used as input into the command-line interface functions of pySCENIC. First, gene regulatory networks were computed using the grnboost2 method in the grn function. Next, enriched motifs were identified using the ctx function, providing the cisTarget v9 databases of regulatory features 500bp upstream, 5kb centered on the TSS, and 10kb centered on the TSS. Finally, individual cells were scored for motifs using the aucell function.

### Identifying over-represented transcription factor motifs in gene lists

The R package RcisTarget^22^ was used to identify enriched transcription factor motifs associated with gene lists, using the cisTarget v9 transcription factor motif annotations and the hg19-tss-centered-10kb-10species.mc9nr database of motif rankings. To compare enrichment between two gene lists, we calculated the difference in normalized enrichment scores for motifs between the two lists and ranked motifs to identify uniquely enriched motifs.

### ATAC-seq sample preparation and analysis

ATAC-seq samples were prepared from OVCA420 cells treated with 10ng/mL of TGFB1 for 0, 1, 3, or 7 days, and the experiment was performed independently twice. Sample preparation was performed as described by Buenrostro et al.^23^. Briefly, nuclei were extracted from 50,000 cells per sample and chromatin was tagmented using the TDE1 transposase provided in the Nextera DNA Library Preparation Kit (Illumina). While the original protocol recommended 2.5µL of enzyme, we found that optimal tagmentation of these samples required 5µL of enzyme at 37°C for 30 minutes with gentle mixing. Finally, ATAC libraries were amplified and sequenced on a NextSeq500 150-cycle high output run, yielding approximately 50M reads per sample.

Raw reads were aligned to the hg38 build of the human genome using Bowtie2^24^ and peaks were called using MACS2^25^ with the following parameters: -q 0.01 -- nomodel --shift -100 --extsize 200 -B --SPMR --broad. Differential motif accessibility was calculated using the R package chromVAR^26^. Briefly, the summits of peaks from all samples were merged, and expanded to a 250bp window, centered on the summit. Motifs from the human_pwms_v2 list included with the package were mapped to the peaks using the matchMotifs() function and then deviations across samples were computed. Significant deviations in motif accessibility were identified using the differentialDeviations() function.

### Code Availability

All code is available on the Github repository hosted at https://github.com/dpcook/emt_dynamics

### Data Availability

All data is in the process of being deposited onto NCBI’s Gene Expression Omnibus and will be available shortly. The GEO accession number will be listed at https://github.com/dpcook/emt_dynamics as soon as it is available.

## Acknowledgements

We thank Pascale Robineau-Charette, Dr. Ken Garson, and the rest of the Vanderhyden lab for their helpful discussion and feedback. We acknowledge StemCore Laboratories and the Ottawa Hospital Research Institute Bioinformatics Core Facility for their technical support and assistance with scRNA-seq and ATAC-seq experiments. We thank Dr. Zev Gartner, Chris McGinnis, and David Patterson for kindly providing MULTI-seq reagents and for their technical support. DPC was supported by a CIHR Frederick Banting and Charles Best Doctoral Award. This work was supported by NSERC grant #RGPIN 2018-0653.8

## Author Contributions

DPC and BCV conceived the study and wrote the manuscript. DPC performed all experiments, computational analysis, and interpreted the data.

---

**Extended Data Figure 1.**
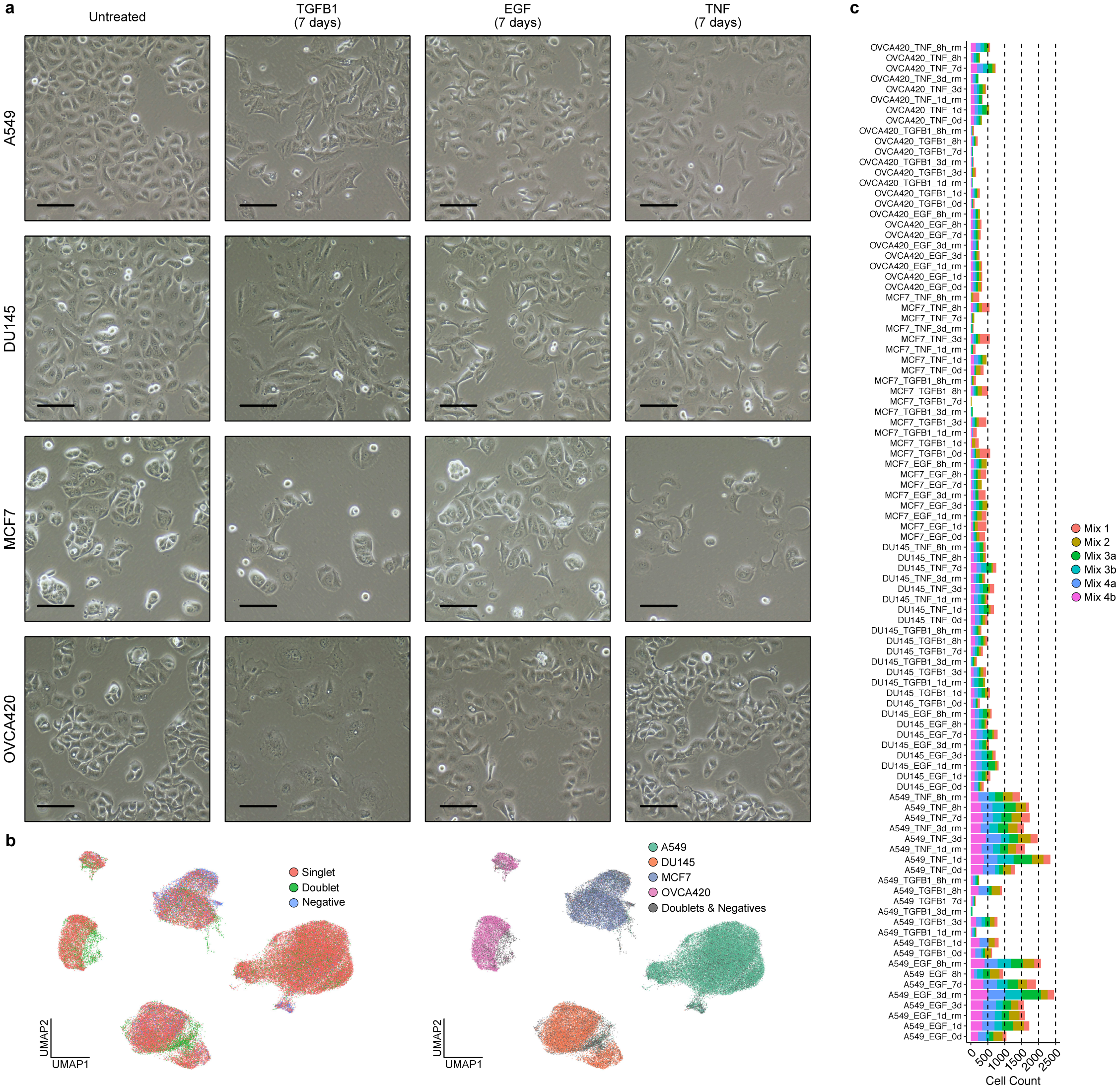
Experimental conditions are associated with EMT-associated morphological changes. **a**, Phase contrast images (20x) of A549, DU145, MCF7, and OVCA420 cells treated with TGFB1, EGF, or TNF. Scale bar = 100µm. **b**, UMAP embeddings of scRNA-seq data from multiplexed EMT time course experiments. Cells are coloured by their annotation after demultiplexing. **c**, Cell counts for each of the 96 samples of the time course experiments after demultiplexing.

**Extended Data Figure 2.**
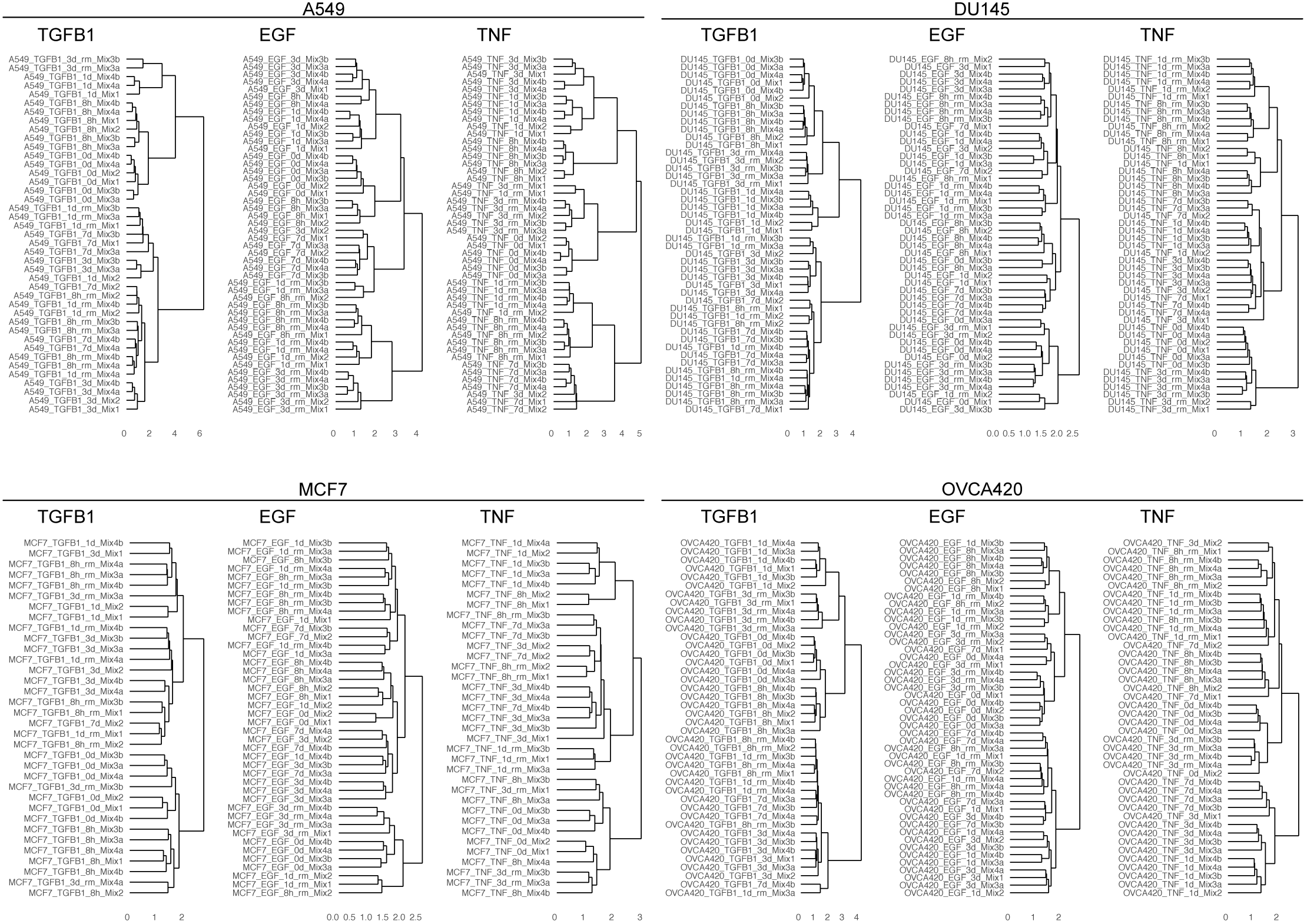
Reproducibility of experimental conditions across replicates. Hierarchical clustering of pairwise correlations (Spearman) from scaled expression values of the top 2000 variable genes of each dataset. Scaled expression data includes a simple linear batch correction.

**Extended Data Figure 3.**
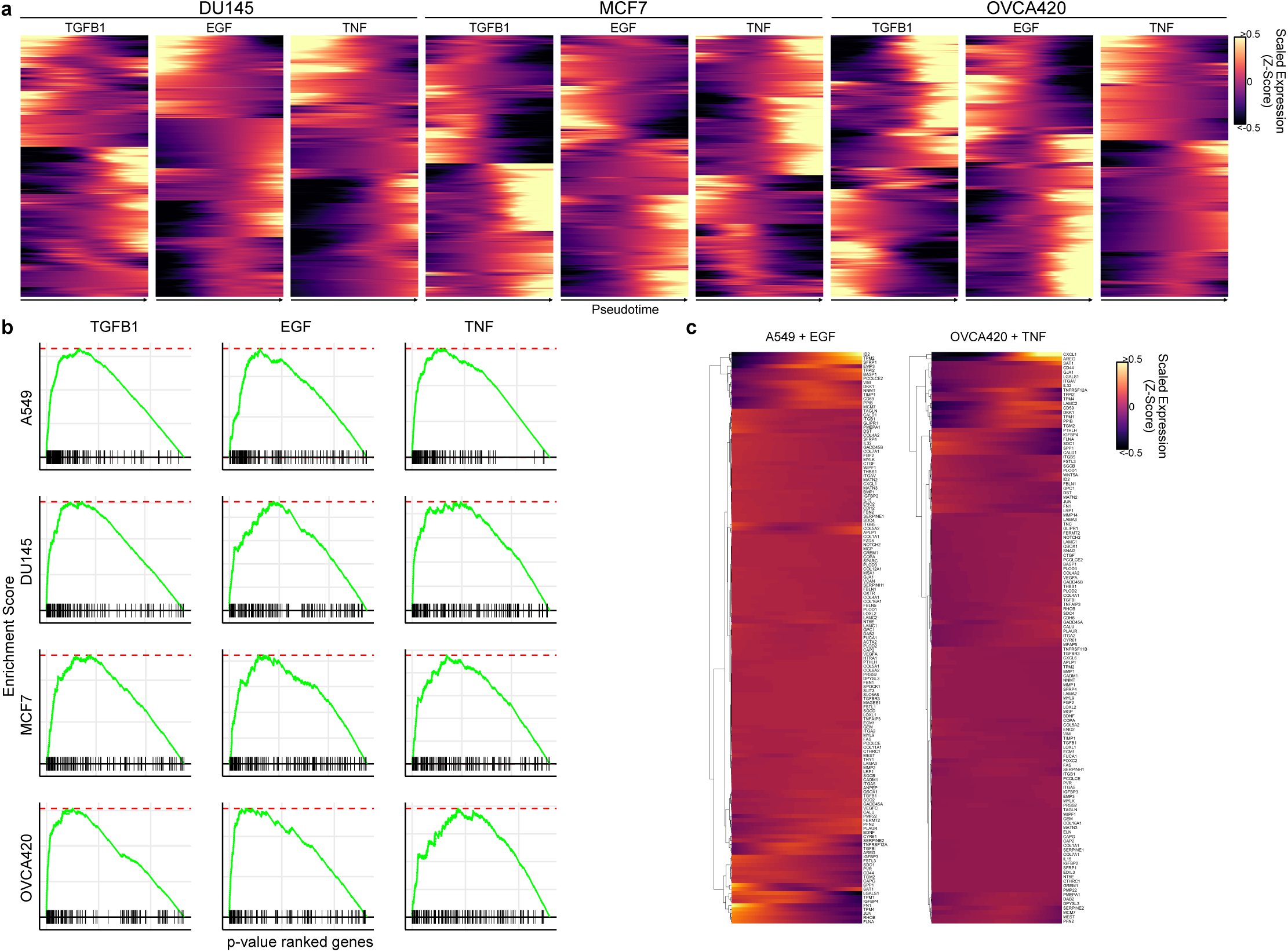
The EMT is associated with diverse transcriptional responses. **a**, Expression dynamics over pseudotime for genes differentially expressed in DU145, MCF7, or OVCA420 time course experiments. **b**, GSEA plots showing the enrichment of the MSigDB EMT hallmark gene set in the most differentially expressed genes for each time course. **c**, Expression dynamics of the MSigDB EMT hallmark gene set throughout pseudotime in A549 cells treated with EGF (left) and OVCA420 cells treated with TNF (right).

**Extended Data Figure 4.**
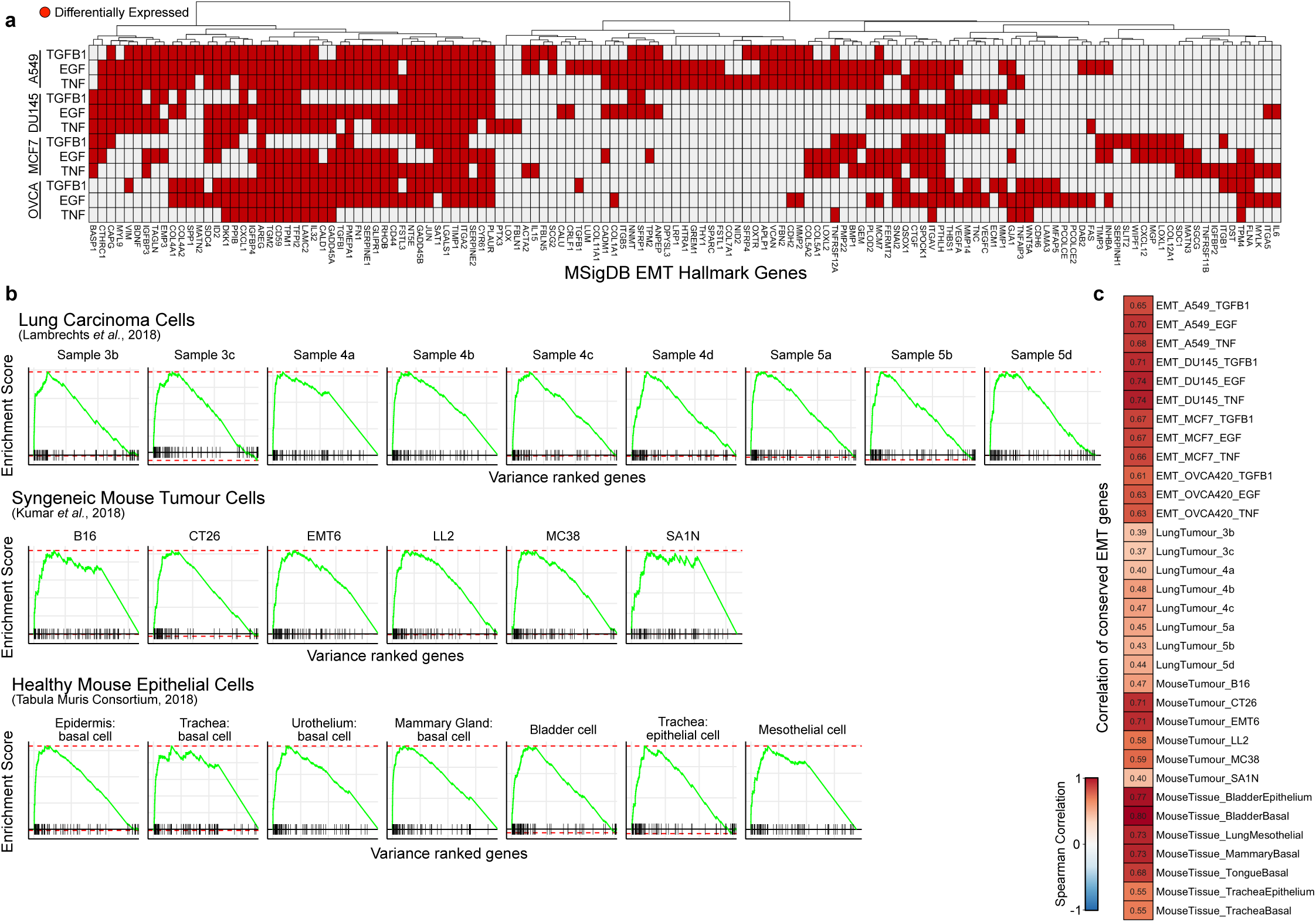
Conservation of EMT gene sets across conditions. **a**, Heatmap showing in which condition each of the MSigDB EMT hallmark genes are differentially expressed (red). The plot only includes genes that are differentially expressed in at least one condition. **b**, GSEA plots of conserved EMT genes in the most variable genes from a variety of epithelial and carcinoma populations. The conserved EMT gene set correspond to the 86 genes that are upregulated in at least 8 time course experiments. Human lung carcinoma scRNA-seq data was acquired from Lambrechts et al.^15^, syngeneic mouse tumour data was acquired from Kumar et al.^16^, and the healthy mouse epithelium data was acquired from the Tabula Muris Consortium^17^. **c**, The average correlation (Spearman) of the same conserved EMT genes and data sets as in (b).

**Extended Data Figure 5.**
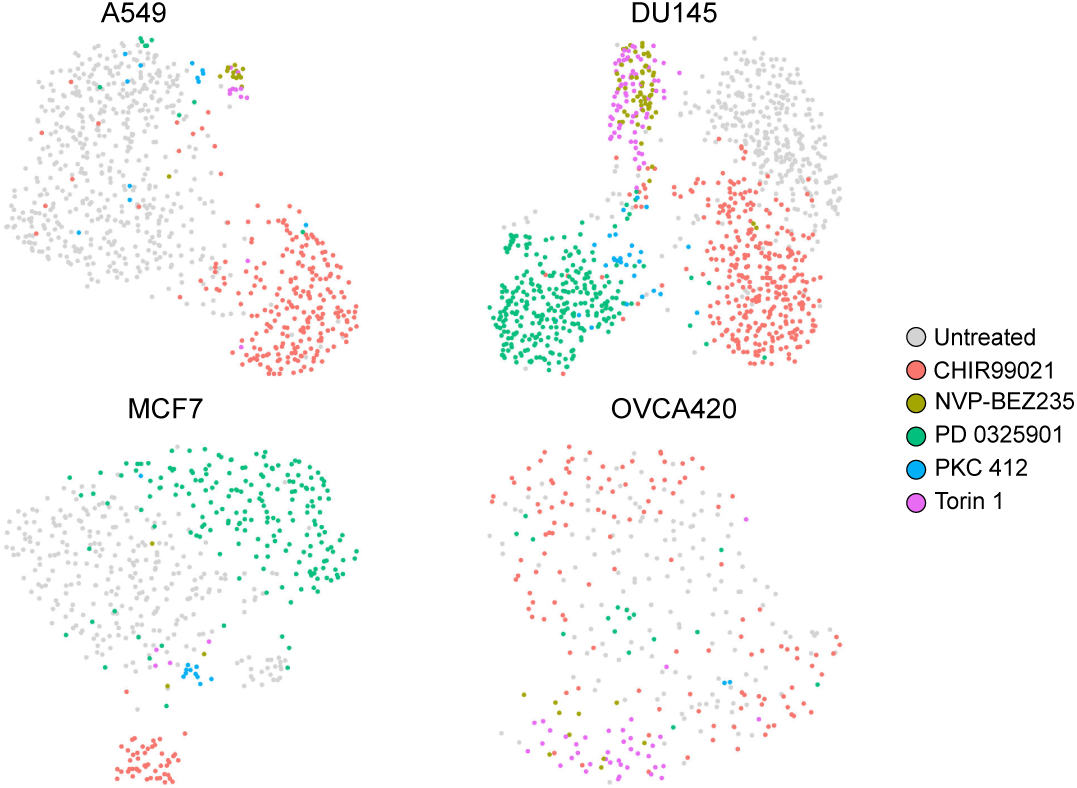
Several kinase inhibitors have large effects on cell state independent of the EMT. UMAP embeddings of untreated cells (grey) and those that had been treated with kinase inhibitors without any EMT induction.

